# Primary motor cortex activity traces distinct trajectories of population dynamics during spontaneous facial motor sequences in mice

**DOI:** 10.1101/2021.02.15.431209

**Authors:** Juan Carlos Boffi, Tristan Wiessalla, Robert Prevedel

## Abstract

We explore the link between on-going neuronal activity at primary motor cortex (M1) and face movement in awake mice. By combining custom-made behavioral sequencing analysis and fast volumetric Ca^2+^-imaging, we simultaneously tracked M1 population activity during many different facial motor sequences. We show that a facial area of M1 displays distinct trajectories of neuronal population dynamics across different spontaneous facial motor sequences, suggesting an underlying population dynamics code.

**Significance statement:** How our brain controls a seemingly limitless diversity of body movements remains largely unknown. Recent research brings new light into this subject by showing that neuronal populations at the primary motor cortex display different dynamics during forelimb reaching movements versus grasping, which suggests that different motor sequences could be associated with distinct motor cortex population dynamics. To explore this possibility, we designed an experimental paradigm for simultaneously tracking the activity of neuronal populations in motor cortex across many different motor sequences. Our results support the concept that distinct population dynamics encode different motor sequences, bringing new insight into the role of motor cortex in sculpting behavior while opening new avenues for future research.

## Introduction

Facial motor control involves descending projections from the primary motor cortex (M1) to the brain stem, through the cortico-bulbar tract, where the facial nerve bilaterally projects to the muscles of the face and neck. Cortical and motor activity involving this tract is highly dynamic and becomes more structured as a consequence of associative learning (1). Associative learning paradigms have extensively contributed to determining the role of M1 in learning new motor skills involving the limbs (cortico-spinal tract) and dexterous movements (2–6). However, relatively little is known about the role of M1 activity outside the context of associative learning and limb movement. Outstanding previous work shows that M1 activity can represent muscle movements (7) and can also encode certain parameters of motor actions such as velocity or direction (8). Both muscle and non-muscle related activity can be detected at M1, which together with its complex network feedback and dynamics makes understanding the functional link between M1 activity and motor behavior non-trivial, even during a single, standardized task (9, 10). Recent research evidences that populations of M1 neurons display markedly different dynamics in reaching versus grasping tasks (11), suggesting that distinct trajectories of M1 population dynamics could arise during different motor sequences. Based on this, we hypothesize that distinct motor actions are encoded by different trajectories of M1 population dynamics. Testing this hypothesis, however, comes with the challenge of recording M1 population activity during multiple different motor sequences. To address this we used a bespoke, fast volumetric 2-photon Ca^2+^ imaging approach based on temporal focusing (sTeFo 2P) (12) to simultaneously sample the activity of M1 neuronal populations across multiple, spontaneous facial motor sequences. We defined, detected and distinguished these spontaneous facial motor sequences from video recordings through a custom algorithm based on behavioral sequencing (13). The combination of these methods revealed that distinct trajectories of population dynamics arise at a facial area of M1 during many different spontaneous facial motor sequences. These distinct trajectories of M1 population dynamics are comprised of transitions between very different population states, and were exclusively observed during specific spontaneous facial motor sequences. Altogether, this supports the concept of a M1 population dynamics code for spontaneous facial motor sequences.

## Results

Spontaneous mouse behaviors have been effectively defined, quantified and studied as sequences of simple behavioral modules that can be systematically detected through image analysis approaches (13). Currently available tools are either designed for tracking body posture (13, 14) or require *a priori* knowledge of specific relevant features for manual training of custom software based on machine learning (15). One prominent algorithm provides effective unsupervised facial movement tracking functionality, producing a multidimensional readout (‘motion masks’) representing certain facial features that change jointly during on-going face behavior (16). However, without previous knowledge of which facial features are relevant for different facial motor sequences, this multidimensional readout must be comprehensively interpreted into a representation of face motor actions, which can be challenging or lead to information loss if dimensionality reduction is used. We in contrast developed an automated classification algorithm to extract behavioral modules from 2-dimensional (2D) videos of the face of head fixed mice (acquired at 60 fps.), and analyzed sequences of these modules to define, detect and distinguish motor sequences as an uni-dimensional readout (Fig. 1a, b). This algorithm relies on the frame by frame categorization of the face conformation via hierarchical clustering of the spatial features in each video frame (Fig. 1b). We vectorize the face conformation displayed in each frame through a histogram of oriented gradients (HOG) transformation (17) and categorize the corresponding HOGs, based on their pairwise similarity. To estimate the number of behavioral modules (different face conformations) present in a video recording, we segmented the smoothed t-distributed stochastic neighbor embedding (tSNE) projection of the HOGs with a watershed transform (14) (see Methods for details). We used this segmentation to split the hierarchical clustering dendrogram of the HOGs into an equal number of branches (clusters). Each cluster of similar face conformations represents one of the aforementioned behavioral modules (‘face categories’). The frame-by-frame sequence of face categories then defines specific motor sequences (Fig. 1b, c, Methods).

**Figure 1:**
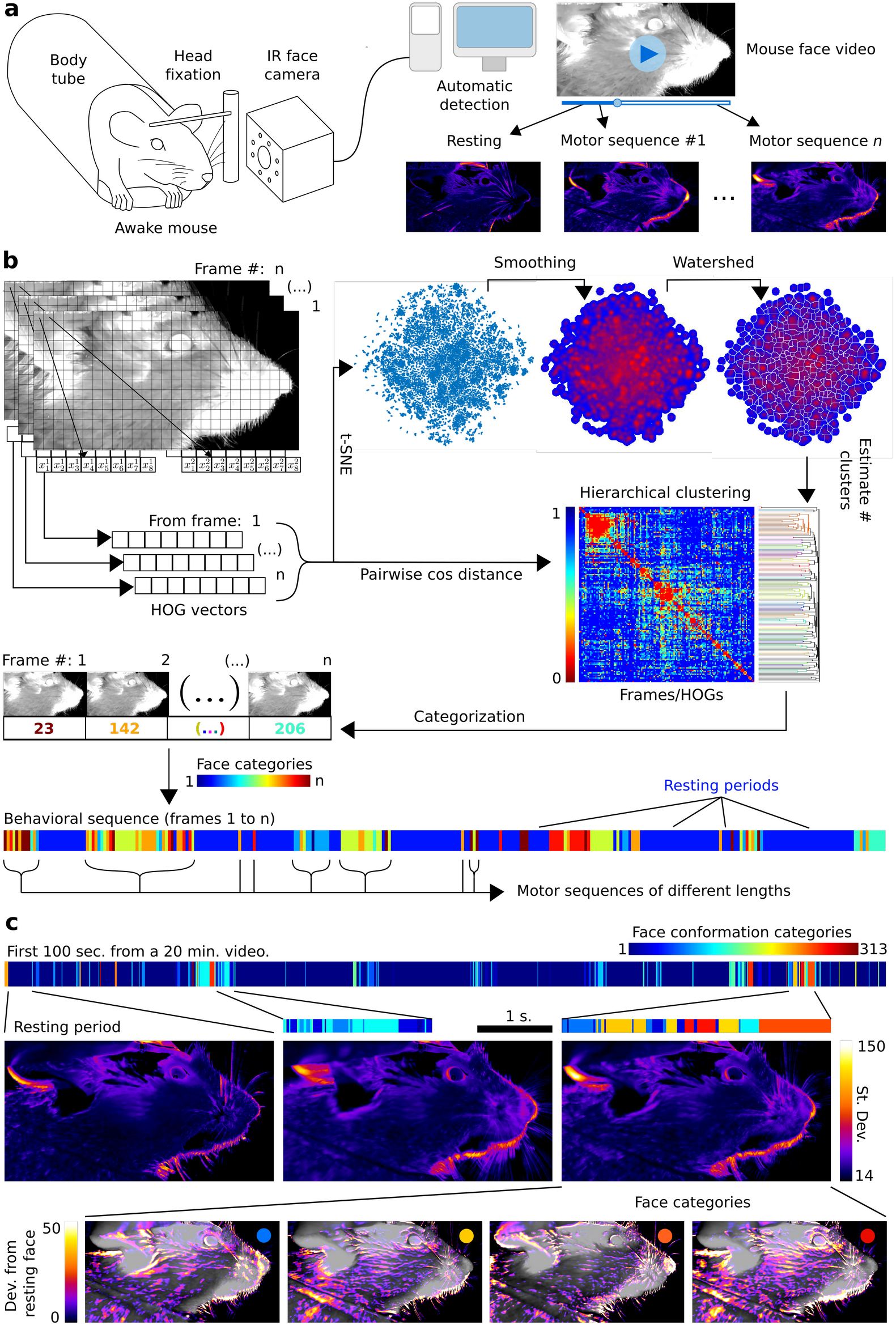
Videographic facial motor sequence interrogation in head fixed awake mice. a) Conceptual diagram of our approach. b) Schematic of the behavioral sequencing algorithm (also see Methods). c) Top: Behavioral sequence obtained from a representative face video (11 experiments, 7 mice). Middle: Example motor sequences and their corresponding standard deviation projections. Bottom: Deviation in gray value from the resting face conformation in identified face categories (indicated by colored dot), highlighting the specific features of each face category.

In all our recordings we observed a single face category that occurred most frequently, covering ~50% of the recording time. We interpreted this as the inactive or resting face conformation. We then delimited motor sequences of different durations as sequences of face categories between resting periods (Fig. 1b, c). In order to explore the datasets for repeated, stereotyped behaviors we evaluated the pairwise Damerau-Levenshtein (DL) distance to estimate the similarity between the detected motor sequences across their different lengths. Only an average of 6% from them showed higher similarity than chance level, suggesting that repetitive or stereotyped motor actions were very infrequent in our recordings. We interpreted the remaining ~44% of recorded motor sequences as different spontaneous motor actions.

We interrogated M1 population dynamics using sTeFo 2P (12) and simultaneously recorded at 4Hz the activity of, on average, 1000 neurons, while tracking facial activity using our algorithm (fig 2a-c). The murine M1 areas associated with face movements are widespread and intermingled with other motor areas (18). In consequence, we focused on a M1 area associated with snout movements, since these include behaviorally relevant motor outputs such as whisking, sniffing, and mouth movements. We deconvolved spike probabilities from the recorded Ca^2+^ signals (fig 2d) to study neuronal population states and their transitions (population dynamics) during the execution of motor sequences and resting periods (fig 2e). Population dynamics traced diverse trajectories during resting periods and motor sequences, which we visualized in principal component space (Fig. 2f). To compare the trajectories of population dynamics without dimensionality reduction, we evaluated the pairwise discrete Frechet distance (FD) as a measure of dissimilarity. Most of the trajectories of population dynamics observed during different motor sequences had lower than chance level FD from the ones recorded during resting states, suggesting that population dynamics at the particular M1 volume imaged (superficial ~450µm^3^) might not be linked to these motor sequences. Nevertheless, we systematically observed up to 11 distinct motor sequences associated with markedly different trajectories of M1 population dynamics. These trajectories displayed higher than chance level FD to all other recorded trajectories, even to those observed during resting periods (Fig. 2g), suggesting a specific association between distinct M1 population dynamics and different spontaneous facial motor sequences.

**Figure 2:**
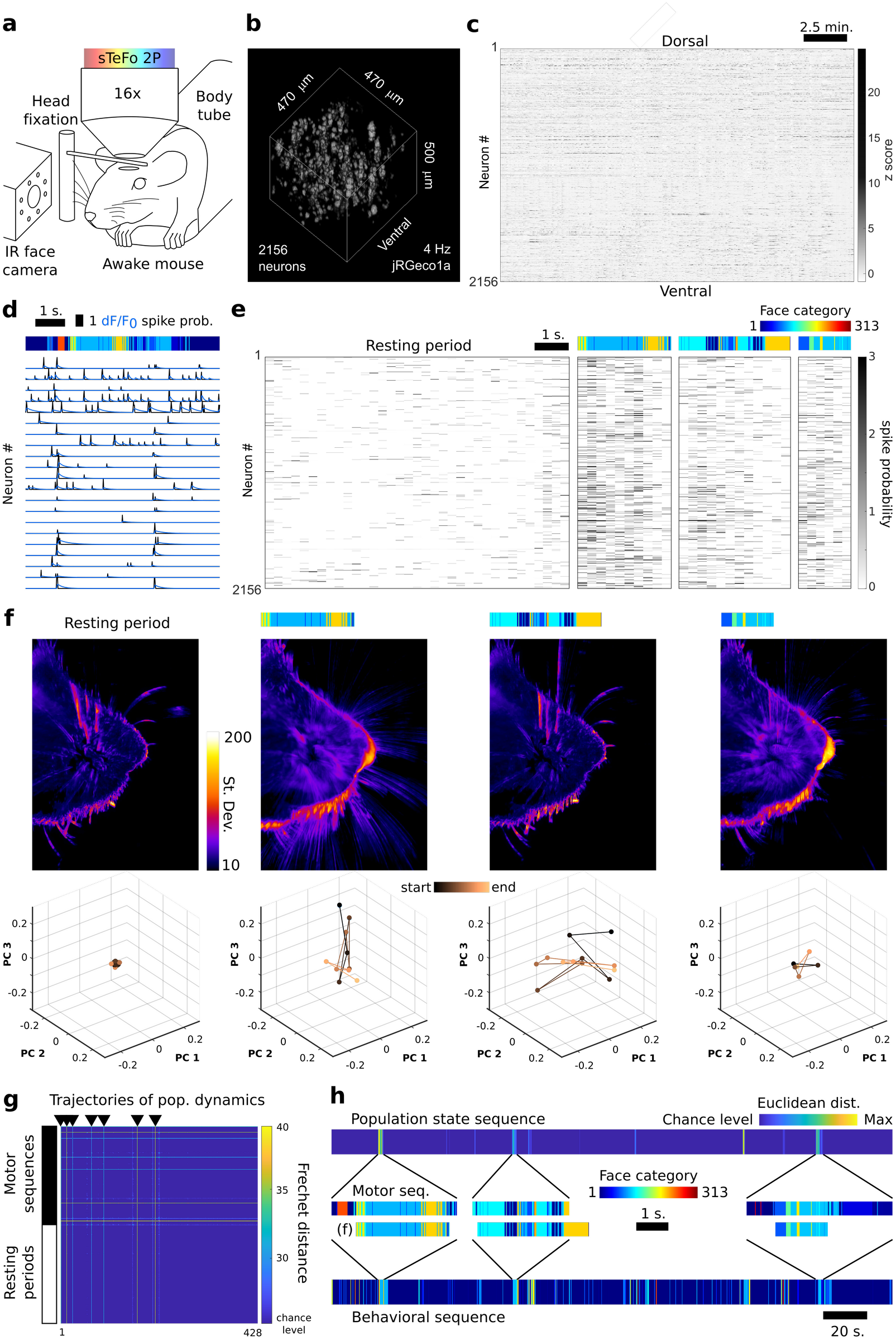
M1 population dynamics during spontaneous face motor sequences. a) Schematic of the experimental setup. b) Volume rendering of the jRGECO1a signal from a representative data set (n = 8 experiments/4 mice). c) Raster plot of the z-scored jRGECO1a signal extracted from all neurons in the dataset shown in (b). d) Face motor sequence with an exemplary subset of associated dF/F_0_ traces (blue) and their corresponding spike probability traces (black). e) Recorded M1 population states transitioned during the corresponding representative face motor sequences, extracted from the dataset shown in (b). f) Standard deviation projections of the gray values in the frames spanning the resting period and three face motor sequences shown in (e), with their corresponding M1 population activity trajectories plotted in low-dimensional principal component space. g) Representative pairwise Frechet distance matrix for trajectories of M1 population dynamics recorded during different face motor sequences (black bar) or resting periods (white bar). Black arrowheads mark trajectories of M1 dynamics with higher than chance level FD to all others (7 in total). h) Top: Euclidean distance between consecutive population state vectors, showing epochs of higher than chance level distance. Middle/Bottom: Corresponding motor sequences delimited by some example epochs with the matching sequences delimited by the behavioral sequencing analysis from panel (f).

If different motor sequences are indeed associated with distinct population dynamics, then these sequences should also be detected through an independent analysis of M1 population dynamics. We therefore evaluated the behavioral sequences delimited by epochs of high Euclidean distance between consecutive M1 population states (Fig. 2h). 83% of the previously detected spontaneous motor sequences (associated to specific trajectories of M1 dynamics) were represented in the motor sequences delimited by high Euclidean distance epochs in population dynamics. This further supports the association between M1 population dynamics and spontaneous facial motor sequences. Finally, we observed similar results when monitoring the face side ipsilateral or contralateral to the brain hemisphere that was imaged, consistent with bilateral cortico-bulbar control of face movements.

## Discussion

Interrogating and interpreting face movements in animal models in an unbiased fashion is challenging, especially when the relevant facial features involved are not fully characterized or are hard to predict beforehand. To overcome this, we developed a bespoke behavioral sequencing algorithm to examine spontaneous facial motor sequences. By combining this approach with simultaneous, volumetric sampling of neuronal population activity, we characterized M1 population dynamics associated with spontaneous face motor sequences. We found a specific association between distinct trajectories of M1 population dynamics and spontaneous face motor sequences. These findings support the concept of a population dynamics code for motor actions, at least for the case of spontaneous face motor sequences. Based on our observations, we hypothesize that the structure of this code would consist of switching population dynamics between a fixed point attractor for the resting period and diverse subspaces or manifolds as flow fields for population dynamics to trace distinct trajectories during different motor sequences (Fig. 2f). Network connectivity within the population would determine the subspace or manifold recruited during a motor sequence. Future research will determine if these subspaces and manifolds are generally low dimensional and overlapping, following the concept of ‘neural modes’ proposed by Gallego and colleagues for the stereotyped, low dimensional behaviors typically used in motor control studies (19). On the other hand, characterizing how contextual inputs (e.g. sensory) switch the population away from and back to resting point attractor dynamics (‘start’ and ‘stop’ signals) is another inspiring future research direction. Altogether, our results and approach bring new insights about the motor control of spontaneous face movement and open new possibilities to study motor systems in health and disease, across conditions such as facial palsy, oro-facial chorea and tic disorders like Tourette’s, and outside the context of the cortico-spinal tract and associative learning tasks.

## Acknowledgements

We acknowledge Lina Streich and Ling Wang for their contribution to microscope assembly and setup, the mechanical and electronics workshops and the laboratory animal resource facility of EMBL Heidelberg for technical assistance. We also thank Hiroki Asari, Cornelius Gross, Santiago Rompani, Marcela Lipovsek and Marina Sabbadini for helpful comments on the manuscript. JCB acknowledges supporting fellowships from the EMBL Interdisciplinary Postdoc (EIPOD) Programme under Marie Skłodowska Curie Cofund Actions MSCA-COFUND-FP (664726). This work was funded by the European Molecular Biology Laboratory (EMBL) and the Deutsche Forschungsgemeinschaft (DFG, German Research Foundation) - project 425902099 awarded to RP.

## Data availability

Data and code are available from the authors upon request.

## Contributions

JCB, TW and RP designed research, analyzed and interpreted data and commented on the manuscript. RP designed the custom 2P microscope and JCB assembled it. TW wrote the video frame categorization code and JCB wrote the behavioral sequencing and Ca^2+^ imaging data analysis pipelines. JCB performed stereotaxic injection and cranial window implantation surgeries and *in vivo* imaging experiments. JCB and TW produced the figures. JCB and RP wrote the manuscript.

## Materials and Methods

### Animals and ethics statement

This work complies with the European Communities Council Directive (2010/63/EU) to minimize animal pain and discomfort. Experimental procedures were approved by EMBL’s committee for animal welfare and institutional animal care and use (IACUC), under protocol number RP170001. 8-16 week old C57Bl6/j mice from the EMBL Heidelberg core colonies were used for experiments, housed in groups of 1-5 in makrolon type 2L cages on ventilated racks at room temperature and 50% humidity with a 12 hr light cycle. Food and water was available ad libitum.

### Surgeries

7-8 week old mice of either sex were used for cranial window surgeries. Animals were anesthetized with a mixture of 40 µl fentanyl (1 mg/ml; Janssen), 160 µl midazolam (5 mg/ml; Hameln) and 60 µl medetomidin (1 mg/ml; Pfizer), dosed in 3.1 µl/g body weight and injected i.p.. After loss of pain reflexes, the fur over the scalp was removed with hair removal cream, eye ointment was applied (Bepanthen, Bayer) and 1% xylocain (AstraZeneca) was injected under the scalp as preincisional local anesthesia. The mouse was then placed in a stereotaxic apparatus (David Kopf Instruments, model 963) equipped with a heating pad (37°C) to preserve body temperature. The dorsal cranium was exposed by removing the scalp and periosteum with fine forceps and scissors to prepare the mouse for M1 ablation or cranial window implantation surgeries. For post surgical care, mice received pain relief (Metacam, Boehringer Ingelheim) and antibiotic (Baytril, Bayer) s.c. injections (0.1 and 0.5 mg/ml respectively, dosed 10μl/g body weight).

### Stereotaxic viral vector delivery and cranial window implantation

For Ca^2+^ indicator expression at M1, recombinant AAV vectors (rAAVs, serotype 1) encoding jRGECO1a under the control of the synapsin promoter (Addgene #100854-AAV1) was stereotaxically delivered as follows. After the dorsal cranium was exposed, a 4mm diameter circular craniectomy was made over M1 using a dental drill (Microtorque, Harvard Apparatus), centered at 1.75 mm anterior and 0.5mm lateral to Bregma. Damage to the dura and bleeding was carefully avoided. rAAV injections were performed at the center of the craniectomy using glass pipettes lowered to depths of 300, 400 and 500 μm, at a rate of ~4ul/ hr using a syringe. ~300nl were injected per spot. After injection, the craniectomy was covered by a round 4mm coverslip (~170μm thick, disinfected with 70% ethanol) with a drop of saline between the glass and the dura. The cranial window was sealed with dental acrylic (Hager Werken Cyano Fast and Paladur acrylic powder) and a head fixation bar was also cemented. The surgical wound was also closed with dental acrylic. Mice were single housed after cranial window implantation and had a recovery period of at least 4 weeks before imaging, for Ca^2+^ indicator expression and for the inflammation associated with this surgery to resolve (20).

### Videographic Behavioral sequencing

We developed a custom library, written in MATLAB code, to perform behavioral sequencing from 2D videography data (https://github.com/prevedel-embl/FaceCat). The videographic data was obtained by imaging one side of the mouse face with an IR sensitive camera equipped with a CMOS OV2710 sensor, IR LEDs for illumination, a 3.6 mm M12 objective (ELP, USBFHD05MT-KL36IR), using a sampling rate of 30 fps at 720p resolution. The library is designed to run without major input by the user, using non-supervised classification algorithms to extract movement bouts from ongoing spontaneous behavior of head-fixed mice.

First, the starting frame of every video is opened in an interactive window that allows the user to specify the area of the image to analyze. HOGs of these areas are calculated as described elsewhere (17). Subsequently the pairwise distances between all HOG vectors are calculated using the cosine distance metric. Unsupervised hierarchical clustering is applied to the resulting distance matrix. To obtain the number of clusters that are present in the data we use a density-based approach following (14). Briefly, the HOG vectors are mapped into two dimensions using t-SNE. A Gaussian kernel is then convolved in both dimensions with the resulting point cloud to obtain a probability density map. Next, a watershed transformation is employed to delineate local maxima in the probability density map. The number of maxima is used as the estimate of the number of clusters present in the data and passed to the hierarchical clustering of the distance matrix. The result is a vector containing one numerical label for every frame of the initial video (classification vector). We defined this classification vector as our behavioral sequence for further analyses.

### In vivo Ca^2+^-imaging of M1 populations

We built a dedicated two-photon microscope based on scanned temporal focusing (sTeFo) which enabled fast volumetric in vivo Ca^2+^ imaging of large M1 neuronal populations. The technical details and working principle is extensively described elsewhere (12), however for our study we utilized a higher repetition laser (10 MHz, FemtoTrain, Spectra Physics) and operated the microscope with the following parameters: The scanning laser power was kept below 191 mW after the objective to avoid excessive heating of brain tissue (12). Field of views of 470 μm^2^ were imaged at 128 px^2^ resolution and spaced in 15 μm z steps in the axial direction to cover 585 μm in depth (39 z steps) at a volume rate of 3.99 Hz. Frames acquired during objective flyback were discarded. Typically we could detect Ca^2+^ signals with good signal-to-noise up to a depth of 500 μm from the pial surface. Mice were briefly (< 1 min) anesthetized with 5% isoflurane in O_2_ for quick head fixation at a custom built stage, with their bodies inside a 5 cm acrylic tube, under our custom two-photon microscope. After head fixation mice fully recovered from the isoflurane anesthesia in less than a minute, showing a clear blinking reflex, whisking and sniffing behaviors and normal body posture and movements. Nevertheless, we waited at least 5 minutes before starting experiments to ensure full recovery from the brief anesthesia. Prior to Ca^2+^-imaging experiments in the awake condition, a pilot group of mice were habituated to the head fixed condition in daily 20 min sessions for 3 days. We note, however, that we did not observe a marked contrast in the behavior of habituated versus unhabituated mice under the microscope during these relatively short 20 min sessions. Thus, imaging sessions typically lasted 20 min, after which the mouse was returned to its home cage. Typically mice were imaged a total of 4 times (sessions), once a week.

### Ca^2+^-imaging analysis

Volumetric imaging data was visualized and rendered using FIJI (21). Motion correction of *in vivo* awake recordings was performed using NoRMCorre (22) on each imaging plane in the volumetric datasets. jRGECO1a signal was extracted, filtered and neuropil corrected from each individual imaging plane using CaImAn (23). Spike probability was estimated using MLSpike (24).

### Statistical analyses

All analyses were performed using MATLAB functions and custom scripts. Dimensionality reduction through PCA, for data visualization purposes only, was performed on motor behavior and resting period associated population states altogether such that their trajectories could be visualized in the same PC space. Discrete Frechet distance (FD) was calculated using the algorithm outlined by Zachary Danziger (2021, Discrete Frechet Distance (https://www.mathworks.com/matlabcentral/fileexchange/31922-discrete-frechet-distance). For determining chance level DL distance and FD we performed 10.000 iterations of resampling with restitution from the recorded DL distances or the FDs recorded dirong resting periods respectively. Chance levels were defined as the 1% quantile from the randomized DL distance distribution and the 99% quantile of the randomized FD distribution. For hypothesis testing, a p value < 0.05 was considered significant.

## References

1. T. Komiyama, et al., Learning-related fine-scale specificity imaged in motor cortex circuits of behaving mice. Nature 464, 1182–1186 (2010).

2. R. Kawai, et al., Motor Cortex Is Required for Learning but Not for Executing a Motor Skill. Neuron 86, 800–812 (2015).

3. A. J. Peters, H. Liu, T. Komiyama, Learning in the Rodent Motor Cortex. Annu. Rev. Neurosci. 40, 77–97 (2017).

4. A. J. Castro, The Effects of Cortical Ablations on Digital Use in the Rat. Brain Res. 37, 173–185 (1972).

5. A. M. Travis, C. M. Woolsey, Motor performance of monkeys after bilateral partial and total cerebral decortications. Am J Phys Med 35, 273–310 (1956).

6. A. J. Peters, S. X. Chen, T. Komiyama, Emergence of reproducible spatiotemporal activity during motor learning. Nature 510, 263–267 (2014).

7. E. V Evarts, Relation of Pyramidal During Tract Activity to Force Exerted Voluntary Movement. J. Neurophysiol. 31, 14–27 (1968).

8. A. P. Georgopoulos, A. B. Schwartz, R. E. Kettner, Neuronal Population Coding of Movement Direction. Science 233, 1416–1419 (1986).

9. A. A. Russo, et al., Motor Cortex Embeds Muscle-like Commands in an Untangled Population Response. Neuron 97, 953–966 (2018).

10. M. Omrani, M. T. Kaufman, N. G. Hatsopoulos, P. D. Cheney, Perspectives on clas-sical controversies about the motor cortex. J. Neurophysiol. 118, 1828–1848 (2017).

11. A. K. Suresh, et al., Neural population dynamics in motor cortex are different for reach and grasp. Elife 9:e58848. (2020). DOI:https://doi.org/10.7554/eLife.58848

12. R. Prevedel, et al., Fast volumetric calcium imaging across multiple cortical layers using sculpted light. Nat. Methods 13, 1021–1028 (2016).

13. A. B. Wiltschko, et al., NeuroResource Mapping Sub-Second Structure in Mouse Behavior. Neuron 88, 1121–1135 (2015).

14. G. J. Berman, D. M. Choi, W. Bialek, J. W. Shaevitz, Mapping the stereotyped behaviour of freely moving fruit flies. J. R. Soc. Interface 11,http://dx.doi.org/10.1098/rsif.2014.0672 (2014).

15. A. Mathis, et al., DeepLabCut: markerless pose estimation of user-defined body parts with deep learning. Nat. Neurosci. 21, 1281–1289 (2018).

16. C. Stringer, et al., Spontaneous behaviors drive multidimensional, brainwide activity. Science 364 eaav7893 (2019). DOI: 10.1126/science.aav7893

17. N. Dolensek, D. A. Gehrlach, A. S. Klein, N. Gogolla, Facial expressions of emotion states and their neuronal correlates in mice. Science 94, 89–94 (2020).

18. K. A. Tennant, et al., The organization of the forelimb representation of the C57BL/6 mouse motor cortex as defined by intracortical microstimulation and cytoarchitecture. Cereb. Cortex 21, 865–876 (2011).

19. J. A. Gallego, M. G. Perich, L. E. Miller, S. A. Solla, Neural Manifolds for the Control of Movement. Neuron 94, 978–984 (2017).

20. A. Holtmaat, et al., Long-term, high-resolution imaging in the mouse neocortex through a chronic cranial window. Nat. Protoc. 4, 1128–1144 (2009).

21. J. Schindelin, et al., Fiji: An open-source platform for biological-image analysis. Nat. Methods 9, 676–682 (2012).

22. E. A. Pnevmatikakis, A. Giovannucci, NoRMCorre: An online algorithm for piecewise rigid motion correction of calcium imaging data. J. Neurosci. Methods 291, 83–94 (2017).

23. A. Giovannucci, et al., CaImAn an open source tool for scalable calcium imaging data analysis. Elife 8, 1–45 (2019).

24. T. Deneux, et al., Accurate spike estimation from noisy calcium signals for ultrafast three-dimensional imaging of large neuronal populations in vivo. Nat. Commun. 7:12682 (2016). DOI: 10.1038/ncomms12682

